# On the rate-limiting dynamics of force development in muscle

**DOI:** 10.1101/2024.08.07.606988

**Authors:** Tim J. van der Zee, Jeremy D. Wong, Arthur D. Kuo

**Affiliations:** Biomedical Engineering Graduate Program, University of Calgary, Calgary, Alberta, Canada; Faculty of Kinesiology, University of Calgary, Calgary, Alberta, Canada

## Abstract

Skeletal muscles produce forces relatively slowly compared to the action potentials that excite them. The dynamics of force production are governed by multiple processes, such as calcium activation, cycling of crossbridges between myofilaments, and contraction against elastic tissues and the body. These processes have been included piecemeal in some muscle models, but not integrated to reveal which are the most rate limiting. We therefore examined their integrative contributions to force development in two conventional types of muscle models—Hill-type and crossbridge. We found that no combination of these processes can self-consistently reproduce classic data such as twitch and tetanus. Rather, additional dynamics are needed following calcium activation and facilitating crossbridge cycling, such as for cooperative myofilament interaction and reconfiguration. We provisionally lump such processes into a simple first-order model of “force facilitation dynamics” that integrate into a crossbridge-type muscle model. The proposed model self-consistently reproduces force development for a range of excitations including twitch and tetanus and electromyography-to-force curves. The model’s step response reveals relatively small timing contributions of calcium activation (3%), crossbridge cycling (3%), and contraction (27%) to overall force development of human quadriceps, with a remainder (67%) explained by force facilitation. The same set of model parameters predicts the change in force magnitude (gain) and timing (phase delay) as a function of excitatory firing rate, or as a function of cyclic contraction frequency. Although experiments are necessary to reveal the dynamics of muscle, integrative models are useful for identifying the main rate-limiting processes.

**Summary statement:** Muscles produce forces relatively slowly, not explained by conventional muscle processes. Quantitative modeling suggests that an intermediate process facilitating force development may be rate limiting.

## Introduction

Skeletal muscle produces force many times slower than its neural excitation. The time course is due to activation dynamics that transform neural excitation into biochemical force-generating capacity, followed by the actual production of force—biophysically attributed to cycling of muscle crossbridges (Fig. 1, center)—against series elastic tissue and the body skeleton. These dynamics are complicated by feedback, in that the resulting body motion also affects the force. The complexity of these interactions obscures which of the preceding processes are rate limiting, and thus most responsible for the relative slowness of force production. This question could be addressed with quantitative muscle models, but there is little agreement or definition for the underlying dynamics. There are presently two main types of models for quantifying force production: (1) mechanistic crossbridge models that focus on the biophysics of force production but not its influence on movements (Huxley, 1957; Zahalak, 1981; Campbell et al., 2018; Newhard et al., 2019; Liu et al., 2024), and (2) phenomenological “Hill-type” muscle models (Hatze, 1977; Hof and Van den Berg, 1981a; Zajac, 1989; van Soest et al., 1993; Thelen, 2003; Mayfield et al., 2023) that interact with tendon and body but lack mechanistic dynamics. There is a need for integration and resolution of both biophysical and phenomenological approaches, to mechanistically determine what limits skeletal muscle force development during movement.

**Fig 1.**
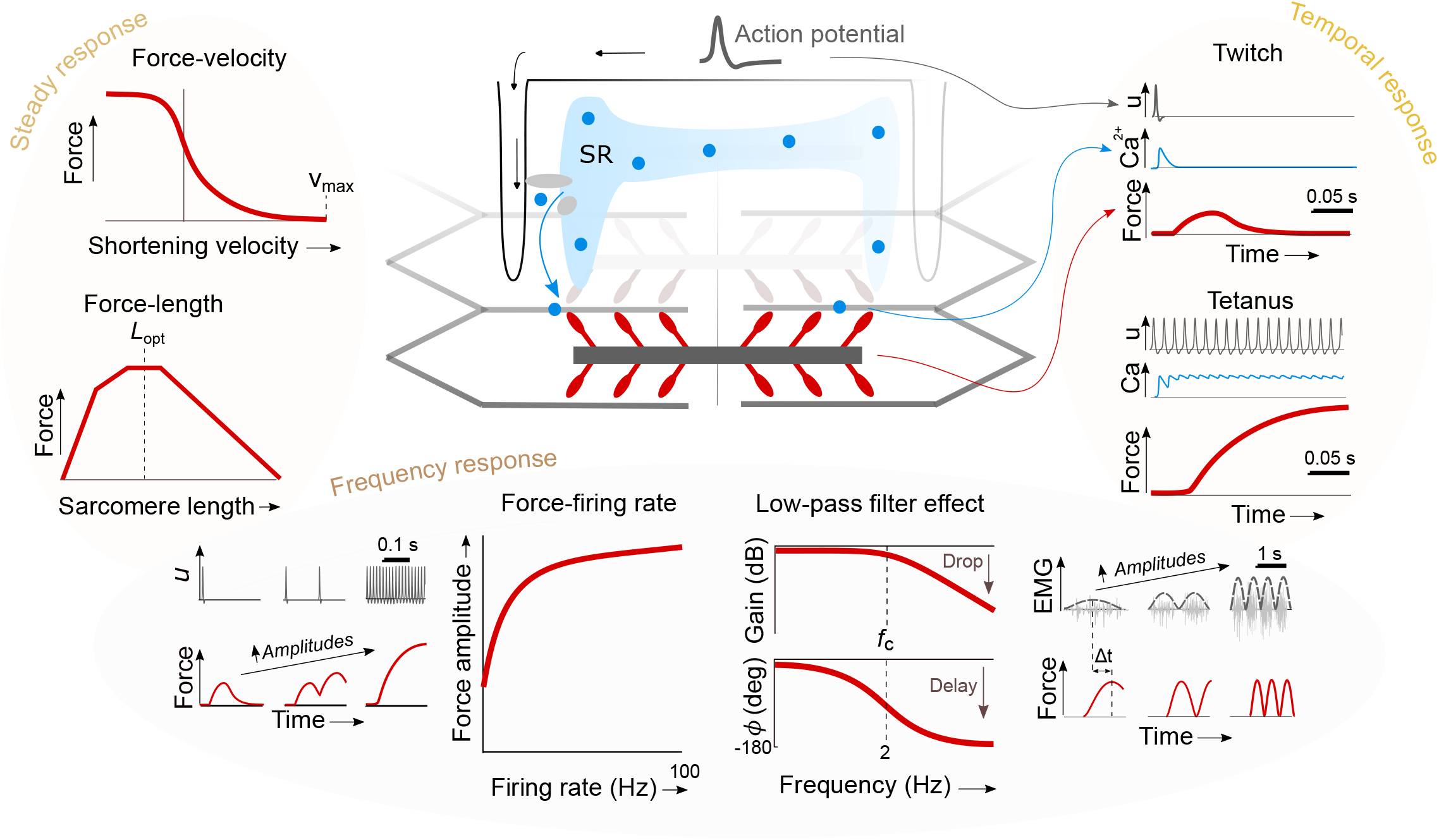
Skeletal muscle and three of its well-known experimental responses to excitation: Steady response, Temporal response, and Frequency response. (Center) Muscle force development dynamics start when an action potential triggering calcium (Ca^2+^) release from the sarcoplasmic reticulum (SR) into the myoplasm, which enables force and work production by crossbridges between thin and thick filaments. (Top left) Steady response include force-velocity (Hill, 1938) and force-length dependencies (Gordon et al., 1966). (Top right) Temporal response includes relative timing of action potential, myoplasmic free Ca^2+^, and force development. Stimuli can range between a single action potential or twitch, and a train of action potentials resulting in tetanus (Hollingworth et al., 1996; Verges et al., 2009). (Bottom) Frequency response refers to dependency of force on excitation frequency. For single fibers or motor units, the force amplitude saturates with increasing action potential frequency or firing rate (Binder-Macleod and McDermond, 1992), yielding a force-firing rate relation. For whole muscle, the magnitude and phase of force exhibits a low-pass filter effect with respect to cyclic contraction frequency, with excitation described by electromyography (EMG; rectified signals shown in inset) (Winter, 2009).

Muscle force development is characterized by several types of experimental observations (Fig. 1). Whereas steady force production is described by well-known force-length (Gordon et al., 1966) and force-velocity (Hill, 1938) relations (Fig. 1, “Steady response”, left), the dynamic effects are evident in time (Fig. 1, “Temporal response”, right) and frequency domains (Fig. 1, “Frequency response”, bottom).

The temporal response shows how excitatory action potentials trigger relatively fast calcium release and binding, followed by relatively slow force development. Frequency response describes how force magnitude and timing vary as a function of excitatory input frequency (Partridge, 1965). For a muscle fiber, the action potential firing rate could be treated as input (Bigland-Ritchie et al., 1983; De Luca and Hostage, 2010) and its force output treated as output, characterized by a force-firing rate curve (Binder-Macleod and McDermond, 1992). For a whole muscle, the periodic electromyographic (EMG) signal could be treated as input (Hof and Van den Berg, 1981b; Zajac, 1989), with the output described by force magnitude (gain) versus the frequency of cyclic muscle contractions (referred to as contraction frequency). The force magnitude (gain) decreases with contraction frequency (Coggshall and Bekey, 1970; Calvert and Chapman, 1977; Crosby, 1978; Genadry et al., 1988; Hof, 1997; Winter, 2009) and its phase lag decreases, similar to a low-pass filter (see Fig. 1, ”Low-pass filter effect”). Although these effects have been well characterized experimentally for decades, it remains unclear how they relate to the underlying biophysical dynamics of muscle.

Force development is described by several types of muscle dynamics (Fig. 2). Musculoskeletal models (e.g., Zajac, 1989) usually distinguish between displacements of contractile and lumped elastic elements (Fig. 2A), between muscle dynamics for activation and contraction, and between body dynamics (Fig. 2B). Muscle dynamics are usually simulated with conventional Hill-type and crossbridge muscle models, which include first-order activation dynamics and contraction dynamics. Activation dynamics transform neural excitation *u*(*t*) into an activation or active state *a*(*t*) Hill-type model contraction dynamics transform active state *a*(*t*) into muscle force *F*(*t*) phenomenologically mediated by the force-length and -velocity dependencies (represented as feedback of muscle-tendon actuator length *L*_MT_ and velocity 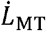, Figs. 2B, 2C). For whole-body movements, it is usually necessary to simultaneously predict both muscle lengths and forces (e.g., *L*_M_ and *F*_CE_ in Fig. 2C), which depend on dynamical interaction between body and series elasticity. Hill-type models typically accomplish this by treating contractile element length as an internal state (Fig. 2B) and using elasticity to relate force and length changes (Zajac, 1989).

**Fig 2.**
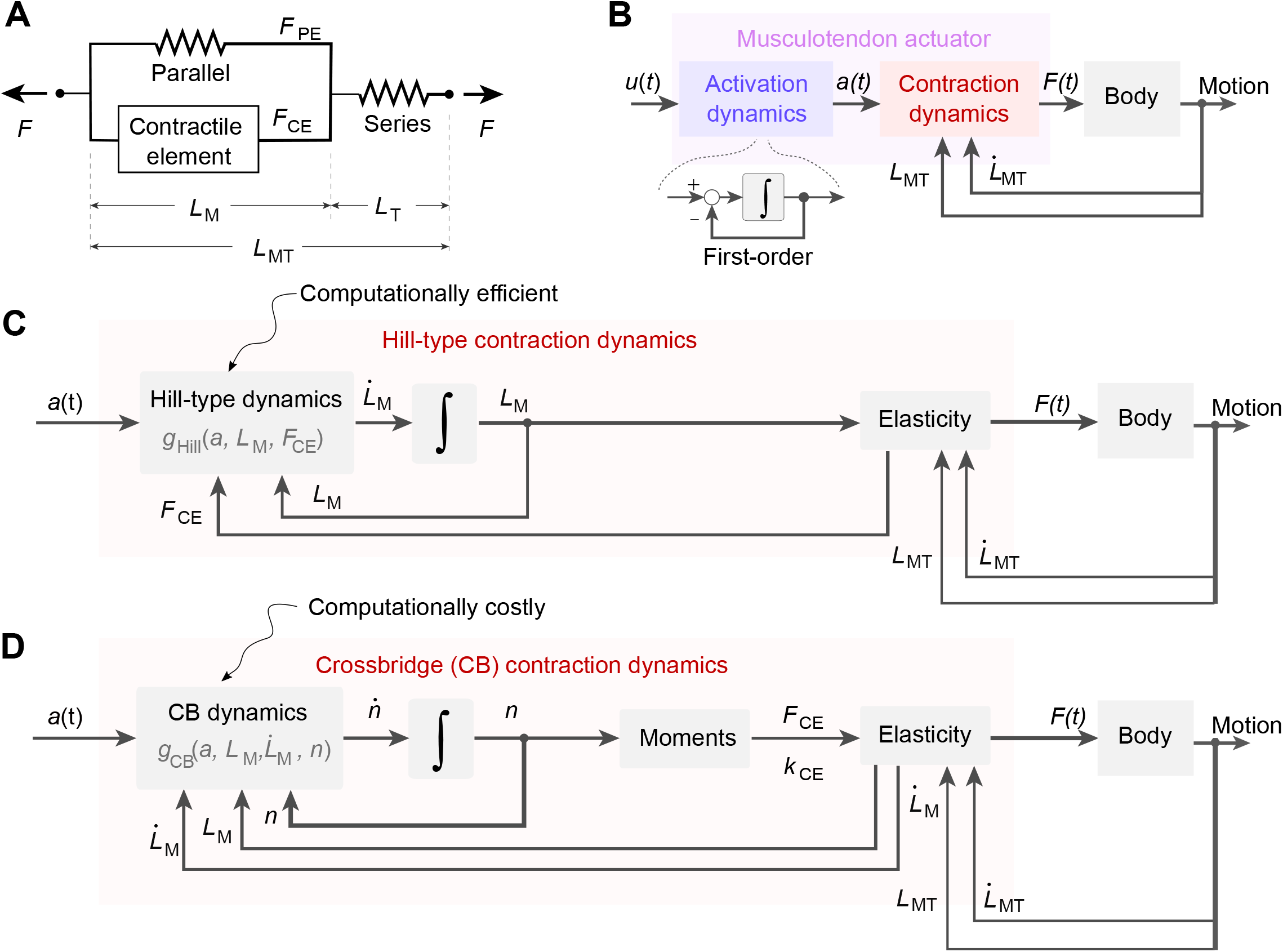
Conventional Hill-type and Huxley-type models of musculotendon actuators. (A) Standard schematic of musculotendon (MT) actuator, featuring muscle contractile element (CE) in parallel with elasticity (PE) and in series with elastic tendon (T). Contractile element, parallel and tendon elements have their own forces, and the tendon element has its own length. (B) Musculoskeletal models feature musculotendon actuators that exert forces on body segments to produce movements, excited by input *u* mediated by activation dynamics to yield activation *a*. Contraction dynamics depend on body motion, incorporated as feedback of muscle-tendon length and velocity (*L*_MT_ and 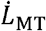). (C) Hill-type contraction dynamics arise from interaction between muscle force-velocity, force-length, and elasticity relations. Series elasticity is used to distinguish contractile length *L*_M_ (treated as a model state) from muscle-tendon *L*_MT_, and parallel elasticity to distinguish contractile force *F*_CE_ from tendon force *F* Note that these dynamics vanish for isometric contraction. (D) Crossbridge contraction dynamics capture distribution of attached crossbridges *n*, with moments that determine contractile force *F*_*CE*_ and stiffness crossbridge binding dynamics that influence force development over time. These models track a strain *k*_CE_. Most such models do not include interactions with body due to high computational cost.

This approach contrasts with more mechanistic models that focus on the dynamics of crossbridge cycling (Fig. 1, center), which may be twice as slow as calcium activation (Zahalak and Motabarzadeh, 1997), and thus at least as important for force development. But most such models cannot be integrated with body and elasticity, except at impractically high computational cost (Fig. 2D; Lemaire et al., 2016; van Soest et al., 2019). The two types of muscle models have different states, dynamics, and parameters, and cannot generally be interchanged within musculoskeletal models. It is also unclear whether either model, or perhaps a combination of the two, can explain force development for whole muscle.

A long-standing inconsistency is how models define muscle activation or active state. Hill (1949) originally treated active state as an inferred force capacity, not directly measurable but estimated from dynamic experiments (e.g., quick-release, Edman, 1970; Edman and Kiessling, 1971; Inbar and Adam, 1976). Others have provisionally defined activation mechanistically in terms of cellular calcium concentrations (e.g., bound to troponin, CaTrop) (Ebashi and Endo, 1968; Hatze, 1977; Hatze, 1978; Zahalak and Ma, 1990) but lacking direct measurements for such concentrations. Most models have therefore defined activation more loosely, as first-order activation dynamics that, when scaled by the steady force-length and -velocity relations (Fig. 1, left) and interfaced with contraction dynamics, yield the muscle force output (Zajac, 1989; Sandercock and Heckman, 1997; Perreault et al., 2003). Associated activation (and deactivation) time constants are determined ad hoc in some models (Zajac, 1989; Thelen, 2003; Millard et al., 2013), and tuned to match overall force development in others (Curtin et al., 1998; van Zandwijk et al., 1998; Lemaire et al., 2016). Consequently, estimated time constants differ considerably, up to ten-fold between definitions (Calvert and Chapman, 1977), and several-fold between similar-seeming Hill-type models (e.g., van Zandwijk et al., 1998; Millard et al., 2013). Another approach is to fit a muscle’s electromyography-to-force transfer function (Calvert and Chapman, 1977; Lloyd and Besier, 2003; Buchanan et al., 2004; Lee et al., 2011; Lee et al., 2013). A transfer function allows for more complex dynamics to non-parametrically describe force production across a range of excitation frequencies, but is also phenomenological and cannot indicate where the dynamics occur. Whereas the early mechanistic approaches to activation (e.g., Hatze, 1977; Hatze, 1978) have lacked data, the more empirical approaches have lacked mechanisms.

However, greater mechanistic insight is enabled by more recent studies. Fast-acting calcium dyes have enabled higher temporal resolution of calcium binding dynamics (Zhao et al., 1996), which play a substantial role in energy expenditure (Barclay et al., 2007). Other studies have revealed additional processes in force production (Irving, 2017). Whereas calcium activation (e.g., CaTrop) was previously thought to initiate crossbridge cycling, newer studies find significant intermediate dynamics, such as for movement of tropomyosin (Kress et al., 1986; Walcott, 2014) that dynamically affects crossbridge binding site availability on the thin filament (McKillop and Geeves, 1993; Campbell, 2014), and for refolding of myosin heads out of a “super-relaxed” state to enable binding to actin (Reconditi et al., 2011; McNamara et al., 2014; Linari et al., 2015; Campbell et al., 2018; Nag and Trivedi, 2021; Liu et al., 2024). These processes can take significant time (e.g., 20-50 ms, Kress et al., 1986; Reconditi et al., 2011), perhaps exceeding that for calcium release and binding to troponin (Hollingworth et al., 1996; Baylor and Hollingworth, 2003). Such dynamics have primarily been studied at the cellular and molecular level thus far (Irving, 2017; Campbell et al., 2018), and could be rate-limiting for whole muscles as well.

Here, the primary objective is to quantify the rate-limiting dynamics of muscle force development. Our approach begins (Part I) by examining prior experimental data for calcium activation and force development dynamics. The data superficially appear to follow first-order processes, but closer examination will show that the activation dynamics seem to vary based on the stimulus train. This suggests the need for quantitative modeling to test whether a more self-consistent set of dynamics can explain such data. We therefore (Part II) apply conventional types of muscle models that have been employed in movement simulation to these data, and show that no conventional model can self-consistently explain muscle force development well, because a new set of dynamics are needed. Finally (Part III), we propose an integrative muscle model to rectify this disparity. We integrate calcium activation with two types of contraction dynamics combining simplified crossbridge dynamics with a Hill-type interface to tendon and body dynamics. To this we add a new set of “force facilitation” dynamics, derived from Part II. The model is tested with a single set of self-consistent parameters, and will be shown to reproduce all three categories of empirical muscle responses (Fig. 1). The model shows that the proposed force facilitation dynamics are necessary to explain relatively slow muscle force development.

### I. Dynamics of biological muscle: Fast activation and slow force

Biological muscle exhibits distinct patterns of calcium activation and force development (Fig. 3). Unavailable at the time of Hill (1949), more recent simultaneous recordings of calcium transients and isometric force development reveal that both resemble first-order processes, but with markedly different time constants. Calcium imaging studies (employing fluorescent dyes in vitro) show that calcium transport and binding to troponin (CaTrop) is relatively fast (Baylor et al., 1983; Zhao et al., 1996), suggesting that other processes are necessary to explain slow force development. These data present an opportunity to test muscle activation models more quantitatively than previously possible.

**Fig 3.**
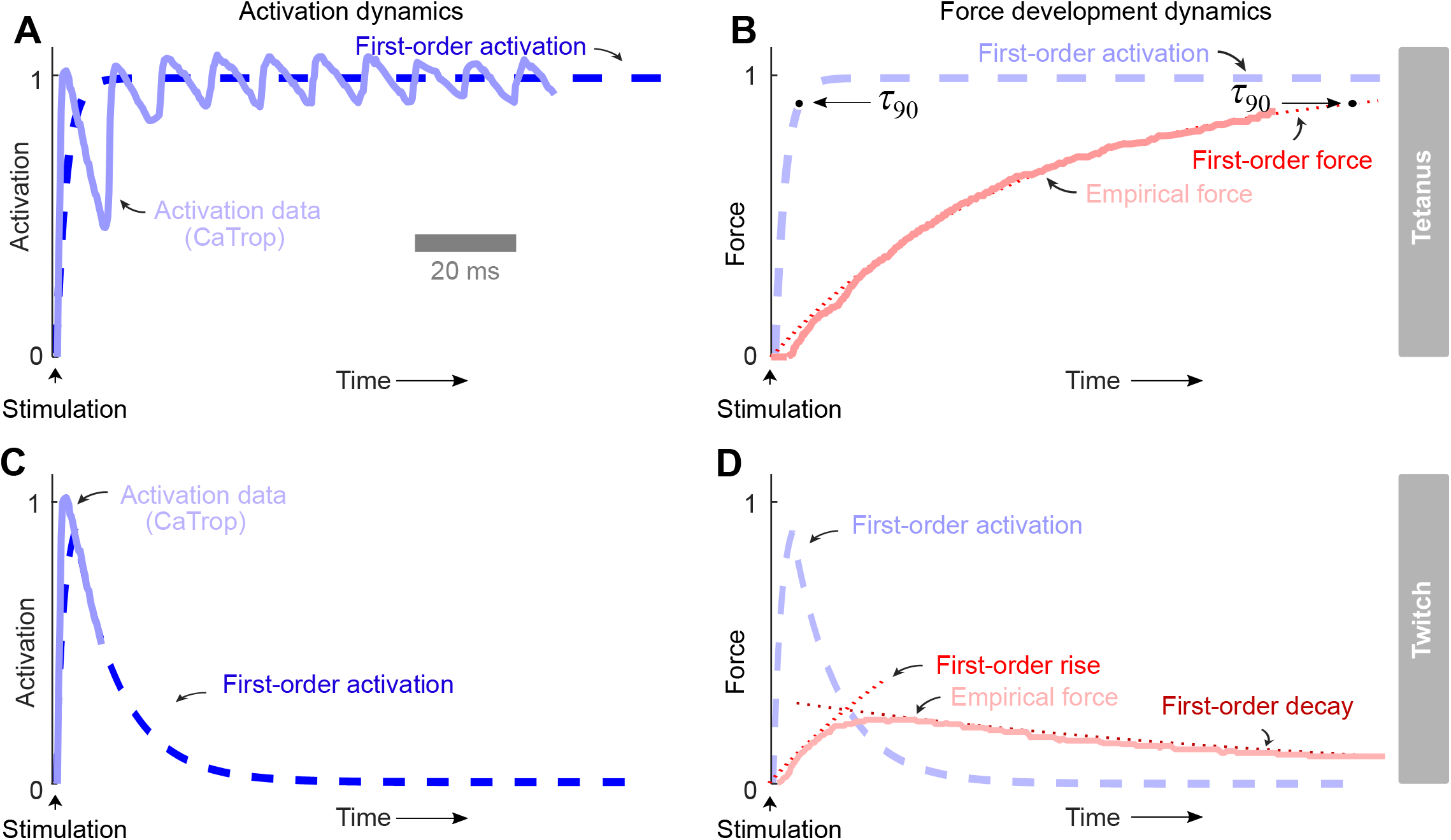
Experimental data for calcium activation and isometric force development dynamics of tetanus and twitch. **A:** Activation has approximately first-order dynamics, describing the time course of calcium bound to troponin (CaTrop; solid line) in response to tetanic electrical stimulation. Calcium activation fluctuates with each action potential, but may be summarized by the step response of first-order dynamics with a relatively fast time constant, quantified with 90% rise time. τ_90_ of about 5 ms (dashed line). **B:** Empirical force (solid light red line) also has approximately first-order dynamics (dotted red line), but about twenty times slower than activation (dashed blue line). **C:** Activation of a twitch also has approximately first-order dynamics (dashed line), with a slower time constant for decay. **D:** Empirical force (solid red line) of a twitch reaches a lower peak value than during tetanus, in line with relatively slow force development dynamics. Force is also well-described by first-order dynamics with different time constants for rise and decay phase (dotted red lines). Data are from mouse extensor digitorum longus muscle fiber bundles at 28C (Hollingworth et al., 1996) (see Supplementary Materials and Methods for details).

Two types of responses are examined here, for step tetanic stimulus (Fig. 3, top row) and single twitch (Fig. 3, bottom row). Tetanic calcium activation (CaTrop from mouse, Hollingworth et al., 1996) is quite fast (Fig. 3A). Ignoring very fast ripples for individual action potentials that do not appreciably affect the force transients (Fig. 3B), the remaining overall time course resembles a saturating exponential, with a 90% rise time (τ_90_, about 2.3 times the exponential time constant) of about 5 ms (Fig. 3A, first-order activation). The resulting force development transient also resembles a saturating exponential but more than twenty times slower, with a τ_90_ of 115 ms (Fig. 3B, first-order force). Additional insights are revealed by the twitch response, which highlights the slower calcium deactivation transient, about six times slower than activation (Fig. 3C). The corresponding twitch force rises and decays much slower than calcium activation (Fig. 3D).

Although these observations all superficially resemble first-order dynamics, they are unlikely to be explained by Hill-type models. Isometric conditions are illustrative because they eliminate Hill-type contraction dynamics, thus isolating first-order activation dynamics as the sole explanation available for force development (e.g., Hatze, 1977; Zajac, 1989; Winters, 1995; Thelen, 2003). It thus appears that Hill-type models should be able to reproduce either fast calcium activation or slow force development, but not both simultaneously. In contrast, crossbridge models have additional dynamics, not present in Hill-type models, that might help account for slower force development. We therefore next examine how both types of conventional muscle models (i.e., Hill-type and crossbridge) compare against these data.

### II. Conventional muscle models: Fast activation *or* slow force

We quantitatively tested whether conventional Hill-type and crossbridge muscle models (Fig. 2) could reproduce empirical activation and force development dynamics. We consulted two data sources: (1) in vitro mouse data for calcium activation and force development as above, and (2) *in vivo* human data of force development (see Supplementary Materials and Methods for details). The latter lack calcium activation estimates, but do include two types of force responses, and thus two ways to test the models. Due to the wide range of model definitions for activation, we considered two extremes for the first-order time constant: a faster and more mechanistic one consistent with calcium-based activation, and a slower and phenomenological one matching force data. Although the data are available in the literature, neither Hill-type nor crossbridge models have previously been tested integratively for both activation and force development. Below, the model dynamics are briefly described and then tested against data.

We implemented both Hill-type and crossbridge models, with some features in common. They share common lumped elements for parallel elasticity, series elasticity (attributed mainly to tendon), and contractile element (Fig. 2A, CE, PE, T respectively; CE and PE share the same length *L*_*M*_). Series elasticity (T in Fig. 2A) is modelled with an exponential-to-linear relationship (Winters, 1995), and parallel elasticity with a quadratic increase beyond its slack length (van Soest and Bobbert, 1993). The active force-length relationship is modeled with a quadratic dependency on length *L*_*M*_ to resemble an inverted U shape (Bohm et al., 2018; Bohm et al., 2019) (see Appendix for parameter values). Both types of models therefore share the same parameter values for these mechanical features.

#### Conventional activation dynamics

We use first-order dynamics (Fig. 2B) between muscle excitation *u* and activation *a*, as previously used in both Hill-type and crossbridge muscle models (Hatze, 1977; Hof and Van den Berg, 1981a; Zajac, 1989; van Soest and Bobbert, 1993; Sandercock and Heckman, 1997; Thelen, 2003; Perreault et al., 2003; Lemaire et al., 2016; Mayfield et al., 2023). We adopted separate time constants for activation and deactivation (i.e., faster forward and slower backward reactions) due to their different underlying mechanisms (i.e., calcium release and uptake respectively, see Appendix for details). As with most existing muscle models, motor unit recruitment and firing rate are lumped into a single quantity *u*, summarizing excitation for the entire muscle.

The two extremes for muscle activation dynamics are as follows. We define “Calcium-based activation” physiologically as the relative amount of calcium bound to troponin regulatory sites (CaTrop) (Ebashi and Endo, 1968; Hatze, 1977; Zahalak and Ma, 1990; van Soest and Bobbert, 1993; Lemaire et al., 2016), based on empirical data (Hollingworth et al., 1996). In contrast, “Force-based activation” refers to a latent and not directly measurable state, selected to approximately agree with force development transients (Zajac, 1989; Sandercock and Heckman, 1997; Umberger et al., 2003; Thelen, 2003; Perreault et al., 2003). Only one model parameter was varied between definitions, i.e. the (forward) activation dynamics time constant. An implication is that a calcium-based activation will reproduce calcium concentration data but not necessarily forces, and a force-based activation the converse.

#### Hill-type contraction dynamics

Hill-type models (Fig. 2C) consider the interaction of muscle force development with series elasticity and body motion (Zajac, 1989). As typical for musculoskeletal models, this is implemented by treating muscle length *L*_*M*_ as a state, using passive elasticity to distinguish from muscle-tendon feedback *L*_MT_. The state derivative is found by inverting the force-velocity relationship (where shortening velocity is − 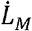). The state equation thus depends on activation *a*, muscle length *L*_*M*_, and contractile element force *F*_CE_:

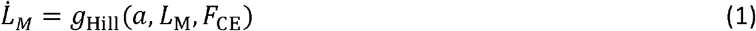

#### Crossbridge cycling dynamics

We implemented a crossbridge model (Fig. 2D) that is compatible with the standard steady muscle behaviors (Fig. 1) and interfaces with body dynamics. Cross-bridge cycling dynamics are dependent on cross-bridge strain *x* and its rate of change 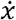, as proposed by Huxley (1957). Cross-bridge strain (rates) are geometrically scaled to whole-muscle fiber *L*_*M*_ and velocity 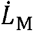 and interfaced with muscle activation *a* as proposed by Lemaire et al. (2016). The distribution of attached crossbridge, 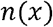 is the state vector, and its state equation depends on muscle activation *a*, muscle length *L*_*M*_, and velocity 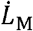 :

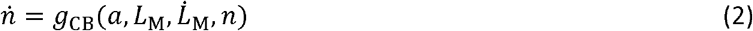

#### Conventional models compared with experimental data

We next examine how these muscle models compare with empirical activation and isometric force. Mouse and human comparisons are presented in separate figures (Figs. 4 and 5), each with both step tetanic excitation and twitch excitation (top and bottom rows, respectively), simulated with two extremes of activation time constants: fast calcium-based and slower force-based (A and B panels, respectively) as subfigures. Within each subfigure, the comparisons include calcium activation (CaTrop) where available, and force development. As with previous modelling studies, twitch and tetanus excitations were simulated using a 5 ms unit pulse (Thelen, 2003; Mayfield et al., 2023) and unit step input respectively (Curtin et al., 1998; Lemaire et al., 2016).

**Fig 4.**
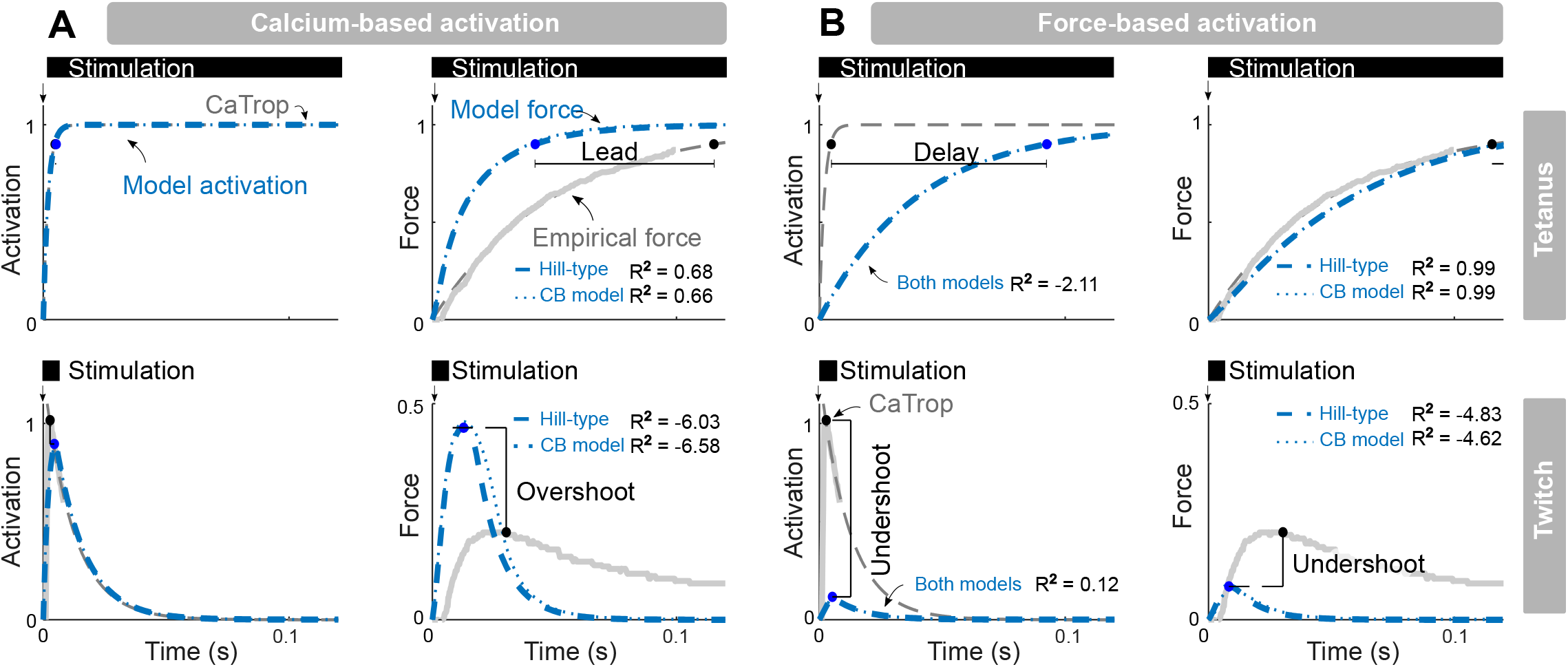
Predictions from conventional muscle models vs. empirical data for temporal response of mouse muscle. Two definitions of activation are compared: **(A)** calcium-based (CaTrop) and a **(B)** force-based latent variable, each with stimuli of step tetanus and single twitch. Calcium-based activation time constant τ_act_ is derived from empirical calcium data (Hollingworth et al., 1996), whereas force-based activation is a latent quantity tuned to match force development data (40 ms vs. 2 ms, respectively). **(A)** Calcium-based definition results in fast muscle activation, but also in overestimation of rate of force development (73 - 76 ms shorter rise time) and twitch force amplitude (119-126% greater) for both Hill-type and crossbridge (CB) models. **(B)** Force-based definition results in slower force development matching data, but also in underestimation of the rate of activation (88 ms longer rise time), twitch force amplitude (62-66% smaller) and twitch activation amplitude (89% smaller). Data are from mouse extensor digitorum longus muscle fiber bundles at 28 C (Hollingworth et al., 1996) ; all quantities are expressed as fraction of the maximal tetanic value (see Supplementary Materials and Methods for details). Model input was square-wave stimulation of indicated duration. Model parameters are the same for force-based vs. calcium-based definitions, except for. τ_act_. Crossbridge model cycling rates were fit on the Hill-type force-velocity relation, yielding: *f*_1_= 287 s^-1^, *g*_1_= 287 s^-1^, *g*_2_= 2049 s^-1^, *g*_3_= 161 s^-1^ (see Appendix for derivation).

**Fig 5.**
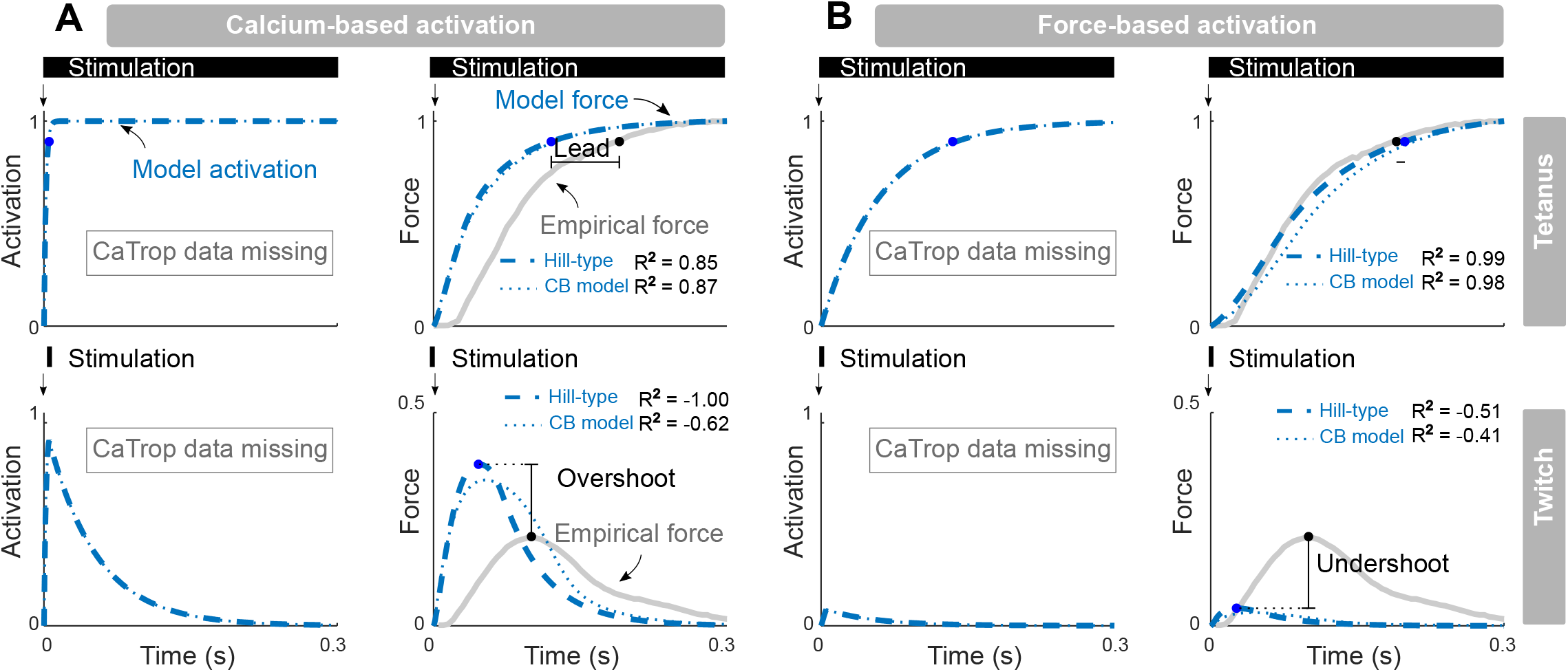
Predictions from conventional muscle models vs. empirical data for temporal response of human quadriceps. Two definitions of activation are compared: **(A)** calcium-based (CaTrop) and a **(B)** force-based latent variable. Two types of stimuli are applied: step tetanus and single twitch. Force-based and calcium-based activation time constants. τ_act_ are 60 ms and 2 ms, respectively. Simulation of model activation is shown, but no CaTrop data were available for human. **(A)** Calcium-based activation results in overestimation of both tetanic rate of force development (65-70 ms shorter rise time) and twitch force amplitude (63-81% greater). **(B)** Force-based activation results in slow force development that can be fitted well to data, but also underestimates twitch force amplitude (80-85% smaller). Data are from human quadriceps muscle *in vivo* (Verges et al., 2009); all quantities are expressed as fraction of the maximal tetanic value (see Supplementary Materials and Methods for details). Model input was square-wave stimulation of indicated duration. All model parameters other than. τ _act_ are the same for both models. Crossbridge model cycling rates were fitted on match the Hill-type force-velocity relation, yielding: *f*_1_= 140 s^-1^, *g*_1_= 140 s^-1^, *g*_2_= 1388 s^-1^, *g*_3_= 78 s^-1^ (see Appendix for derivation).

These comparisons will show that activation dynamics contribute more to force production than other muscle dynamics. This is because (near-) isometric conditions downplay the effect of contraction dynamics and interaction with body, and crossbridge cycling dynamics needed to be relatively fast to comply with force-velocity relations (see Appendix for details). As both models share the same activation dynamics, both Hill-type and crossbridge models produce very similar simulations (dotted vs. dashed blue traces, respectively in Figs. 4 and 5). We therefore describe general “model” results here, only distinguishing between Hill-type and crossbridge models where specifically needed.

We first compare the models with in vitro mouse data from fast-twitch extensor digitorum longus muscle (Fig. 4), including simultaneous recordings of CaTrop and force (Hollingworth et al., 1996) (see Supplementary Materials and Methods for details). As expected, a calcium-based activation model can successfully replicate CaTrop transients for both step tetanic and twitch excitation (Fig. 4A). However, combining this activation with contraction dynamics results in forces far faster than empirical data. Additionally, the models’ twitch force response exhibits far greater amplitude than data show (Fig. 4A). Alternatively, selecting a slower force-based activation time constant enables models to reproduce step tetanic force development (Fig. 4B). However, the associated activation dynamics must then be far too slow to match empirical CaTrop data. Moreover, a model proficient in replicating step tetanic force performs inadequately for twitch force, with far too low an amplitude (Fig. 4B). Consequently, conventional models can predict either activation or force transients observed experimentally, but not both (Fig. 4). Altogether, the contraction dynamics of both Hill-type and crossbridge muscle models prove to be excessively fast to account for these empirical data.

Similar results were obtained for *in vivo* human quadriceps muscle data (Fig. 5), including dynamometer-based estimates of muscle force (Verges et al., 2009) (see Supplementary Materials and Methods for details). Lacking calcium activation data for human, the models are tested by their ability to reproduce force development data for both step tetanic and twitch stimuli. Again, we consider two possibilities for model activation time constants, selected to correspond to (assumed) calcium-based activation or to agree with empirical force data. As expected, calcium-based activation is relatively fast, and the subsequent model contraction dynamics are insufficient to explain the relatively slow forces observed experimentally. The models produced far faster tetanic forces and larger twitch forces compared to empirical data (Fig. 5A). The alternative of slower, force-based activation was able to reproduce step tetanic force response well, but then greatly underestimated twitch force amplitude (Fig. 5B). Altogether, even in the absence of empirical CaTrop activation data, current first-order models of activation dynamics cannot reproduce both twitch and tetanic force responses.

These findings confirm the suspicions from Part I, and show that conventional muscle models cannot simultaneously predict twitch and tetanic force development data. Models with calcium-based activation can reproduce fast calcium activation dynamics (when empirical data available, Fig. 4), but not the observed slow force development, because their contraction dynamics are too fast to reproduce data. Models with force-based activation can reproduce tetanic force data (Figs. 4 and 5), but at the expense of not agreeing with twitch force data (Fig. 5), and/or requiring unrealistically slow calcium activation (Fig. 4). This evidence suggests that muscle exhibits an additional set of dynamics, not explainable by any first-order model of activation dynamics, nor by either type of contraction dynamics.

### III. An integrated muscle model with force facilitation dynamics

We next propose a new muscle model that resolves many of the above issues, called the CaFaXC (after calcium-facilitation-crossbridge-contraction) model (Fig. 6). It integrates several features of existing models, including calcium-based activation dynamics (*c*, representing physiological CaTrop; Fig. 6A) and contraction dynamics that encompass both length- and body feedback and crossbridge cycling dynamics. However, it also inserts a second and slower set of first-order dynamics, termed force facilitation dynamics (Fig. 6B), before contraction dynamics. Studies have revealed force development is affected by processes such as tropomyosin movement (e.g., Moore et al., 2016) and myosin unfolding (e.g. Linari et al., 2015), although it is presently uncertain which of them is rate limiting. Rather than modeling them explicitly, we provisionally model their lumped effect with an additional latent force facilitation state *r* that reflects relatively slow force development. Below we first describe the model and then test it against experimental calcium and force data (Fig. 1).

**Fig 6.**
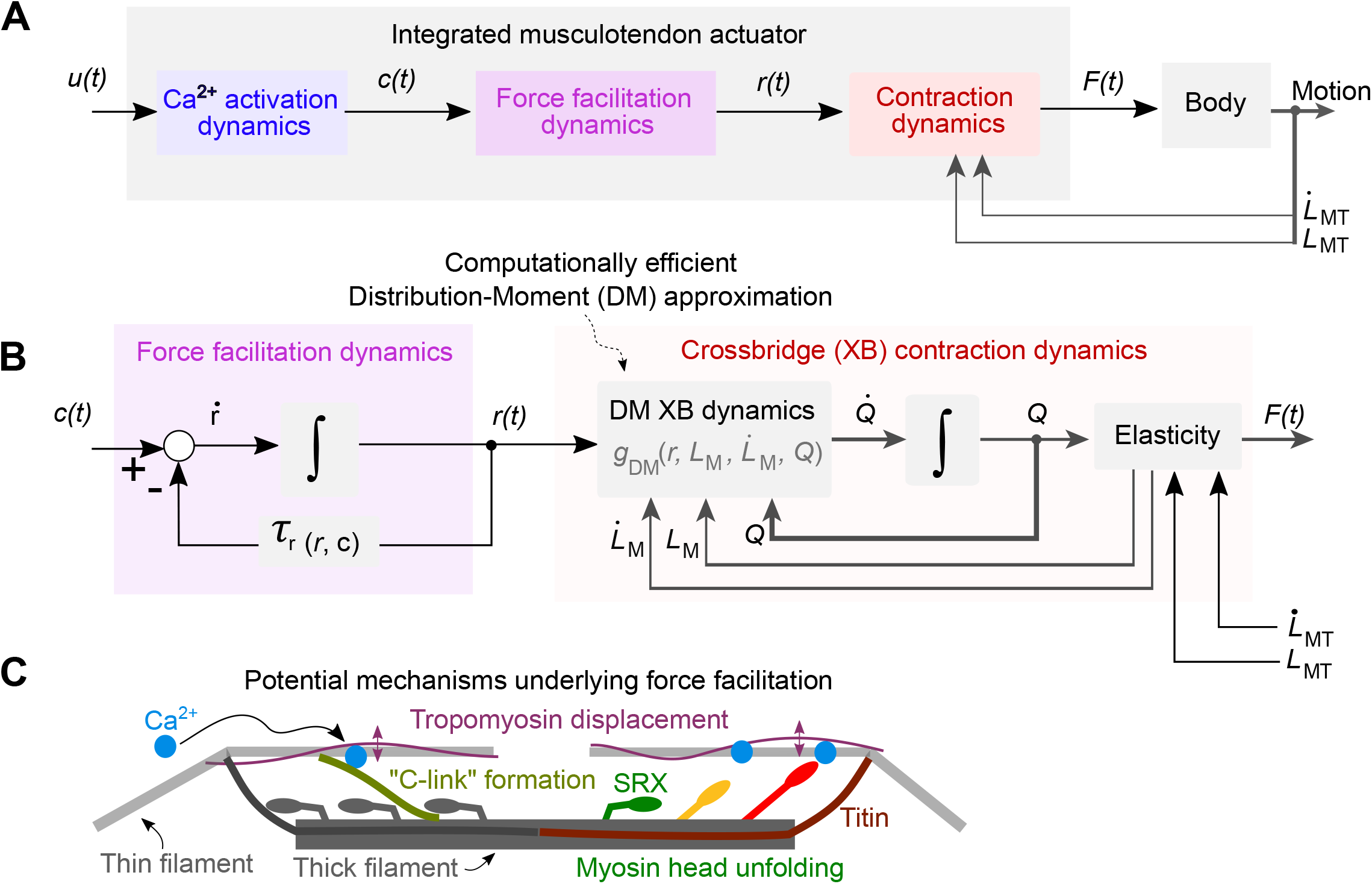
Proposed calcium-facilitation-crossbridge-contraction (CaFaXC) model with force facilitation dynamics. **(A)** Proposed model includes three sets of dynamics: calcium (Ca^2+^) activation dynamics, force facilitation dynamics, and contraction dynamics. Calcium activation dynamics are first order, with state explicitly defined as calcium bound to troponin (CaTrop), referred to as calcium activation *c*. **(B)** This activates a set of first-order force facilitation dynamics with state *r* and time constant. τ_r_ that precede crossbridge contraction dynamics. Crossbridge dynamics are modeled with a Distribution-Moment (DM) approximation. The proposed force facilitation dynamics are critical for allowing calcium activation *c* to match experimental calcium measurements yet also produce force as slowly as experimental force measurements. **(C)** Force facilitation is a simplification of more complex dynamics that are left unmodeled. Potential mechanisms include tropomyosin movement (Moore et al., 2016), myosin head unfolding out of a quiescent “super-relaxed” (SRX) state (Linari et al., 2015; Ma et al., 2023) possibly mediated by titin activation (Squarci et al., 2023), and formation of myosin-binding protein C links between thin and thick filaments (Hessel et al., 2024; Song et al., 2021). A single force facilitation state is used to approximate the lumped effect of such dynamics, with time constant tuned to match force data.

As with the crossbridge model (see Part II), we adopted the approach of Lemaire et al. (2016) to interface crossbridge cycling dynamics with body feedback and muscle length changes. To reduce the computational cost, we also simplified the model using the DM approximation for the continuous distribution (Zahalak, 1981). The same diagram as for conventional crossbridge models (Fig. 2D) applies, but now defining three ordinary differential equations for *Q*_0_ through *Q*_2_, the moments of the distribution:

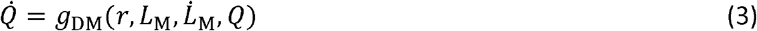

The distribution moments may be regarded as equivalent to contractile element stiffness, *k*_CE_, force *F*_CE_, and elastic energy, or to the distribution’s amplitude, mean, and standard deviation. For body interaction, these moments were combined with elasticity to track muscle length and velocity, including appropriate scaling to ensure compatibility of crossbridge forces and displacements with the whole muscle (see Appendix). Owing to the DM approximation, CaFaXC simulates contractions about 700 times faster than conventional crossbridge models and only about two times slower than Hill-type models (see Appendix).

#### Model parameters

The model’s parameters are determined as follows (see Appendix for details). The calcium-based activation dynamics have activation and deactivation time constants determined either from empirical CaTrop (for comparison with mouse data, described below), or from twitch and tetanic force responses (for human, described below). Force facilitation also has forward and backward time constants, here tuned to match empirical force, similar to force-based activation in Hill-type models. The contraction dynamics require four crossbridge rate parameters, here constrained by the force-velocity relationship similar to described previously (Huxley, 1957; Zahalak, 1981; van Soest et al., 2019). Three of them may be mapped to three Hill-type force-velocity parameters, for example maximal shortening velocity, force-velocity curvature, and eccentric force asymptote (see Appendix). The fourth parametric degree of freedom is a free parameter, unexamined here, that effectively relates shortening velocity to energetics, somewhat like some models that link Hill-type models to phenomenological energetics estimates (e.g., Umberger et al., 2003). The proposed model also has length and force scaling parameters to ensure compatibility between crossbridge strains and muscle length changes (Lemaire et al., 2016). These scaling parameters may be mapped from optimal fiber length and maximum isometric force of Hill-type models (see Appendix). The model’s parameters may therefore largely be mapped from Hill-type models, except for two additional time constants for force facilitation dynamics.

#### Integrated muscle dynamics compared with experimental data

The CaFaXC model is tested against step tetanic and twitch temporal response data (Fig. 7). Starting with mouse tetanic response (Fig. 7A), the fast activation dynamics yield fast activation rise (τ_90_ = 5 ms), in line with empirical CaTrop data (Hollingworth et al., 1996). Using the empirical force development data to determine the force facilitation dynamics, the model also reproduces relatively slow force development (τ_90_ = 102 ms, Fig. 7A). These time constants provide quantitative contributions to overall force development time: calcium activation 5%, force facilitation 77%, crossbridge dynamics 3%, and contraction dynamics 16%. Without modification, the same model reproduces a twitch response that shows fast activation and slow force. Unlike conventional models (Fig. 4B), the integrated model predicts much smaller twitch force amplitudes compared to tetanic force (1:5 ratio, Fig. 7A), similar to experimental data. Both tetanic and twitch responses were dominated by force facilitation dynamics and reproduced data well.

**Fig 7.**
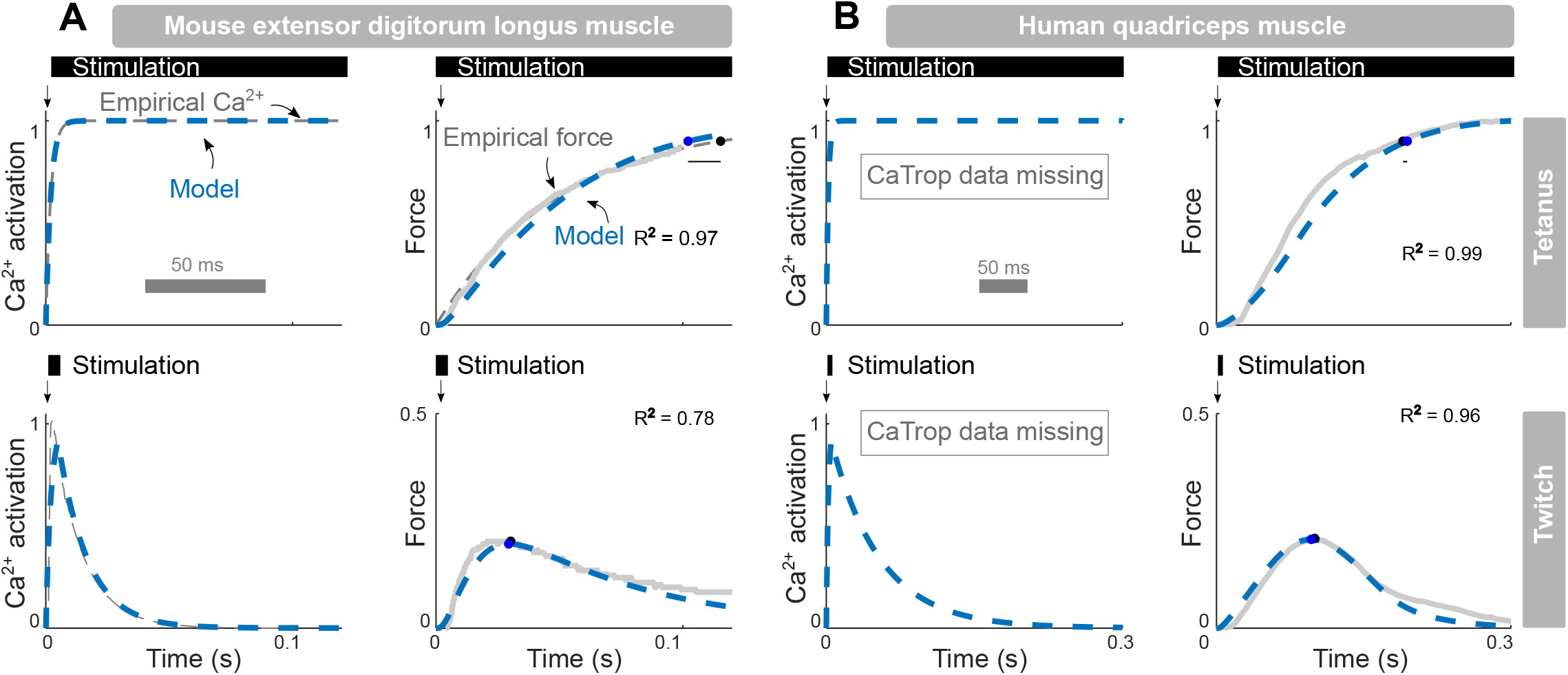
Proposed model predictions vs. empirical data for temporal response of human quadriceps and mouse fast-twitch muscle. Proposed model produces realistic force development in mouse (A) and human muscle (B) and mouse muscle. Model produces slow tetanic force development in human (4 ms longer rise time than empirical) and mouse (13 ms shorter rise time than empirical). For mouse muscle, predictions of activation match empirical estimates in both twitch and tetanus. As there is no data of human muscle activation, activation is only shown for model. Model twitch amplitude matches empirical force amplitude in human (1% smaller than empirical) and mouse (3% smaller than empirical). Mouse data (A) are from extensor digitorum longus muscle fiber bundles *in vitro* at 28 C (Hollingworth et al., 1996) and human data (B) are from quadriceps muscle *in vivo* (Verges et al., 2009) (see Appendix for details). Ca^2+^ activation *c* is defined as the fraction of Ca^2+^ bound to troponin; all quantities are expressed as fraction of the maximal tetanic value. Model input was square-wave stimulation of indicated duration. Mouse parameter values: τ _act_ = 2 ms, τ _deact_ =12 ms, τ _fac_ = 35 ms, τ _def_ = 60 ms,*f*_1_ = 287 s^-1^,*g*_1_ = 287 s^-1^, *g*_2_= 2049 s^-1^, *g*_3_= 161 s^-1^. Human parameter values: τ _act_ = 2 ms, τ _deact_ = 50 ms, =τ _fac_ 60 ms, =τ _def_ 20 ms, *f*_1_= 140 s^-1^, *g*_1_= 140 s^-1^, *g*_2_ = 1388 s^-1^, *g*_3_ = 78 s^-1^ (see Appendix for derivation and definitions).

The model also compares well with human data (Fig. 7B). Lacking human CaTrop measurements comparable to mouse, the model is instead compared against the combination of twitch and tetanus data. With a single set of parameters, the model reproduces both of the relatively slow twitch and tetanic force responses (τ_90_ = 194 ms, Fig. 7B) reasonably well. The contributions were calcium activation 3%, force facilitation 67%, crossbridge dynamics 3%, and contraction dynamics 27%. Again, it produces a small twitch-to-tetanus ratio similar to data (1:5, Fig. 7B), unlike conventional models (Fig. 5B). Although the associated activation dynamics are not presently testable for human, the model predicts relatively fast activation (τ_90_ = 5 ms, Fig. 7B) and slow force facilitation, consistent with mouse data. Apart from different time scales, the model suggests that activation and force development dynamics appear qualitatively similar for mouse and human. A self-consistent model for each can simultaneously reproduce activation (in mouse) and force responses (in mouse and human), including both twitch and tetanus.

We next examine whether a single set of CaFaXC parameters for human quadriceps muscle can reproduce multiple empirical observations. First, since the model’s steady responses were constrained to empirical force-length and force-velocity relations for human quadriceps, these are reproduced relatively well (R^2^ = 0.85 and R^2^ = 0.96 respectively, Fig. 8, left panel). The model also reproduces both twitch and tetanus force development (R^2^ = 0.96 and R^2^ = 0.99 respectively, Fig. 8, upper right panel), with tetanus rise time (._90_ = 194 ms) and twitch amplitude similar to empirical values (Fig. 7). The same model then makes independent predictions of frequency response data, including the EMG-to-force relationship as a function of cyclic contraction frequency (Fig. 8, lower right). Using the same parameter values above, the model shows a decrease in gain and an increase in phase lag between EMG and force, in agreement with empirical data (R^2^ = 0.93 and R^2^ = 0.82 respectively). In addition, the model predicts a saturating increase for the force-firing rate relationship, similar to experimental data (R^2^ = 0.96, Fig. 8 lower right). Altogether, the model self-consistently reproduces multiple muscle responses, including classic steady force-length and velocity relations, temporal responses, and frequency responses, across a range of conditions.

**Fig 8.**
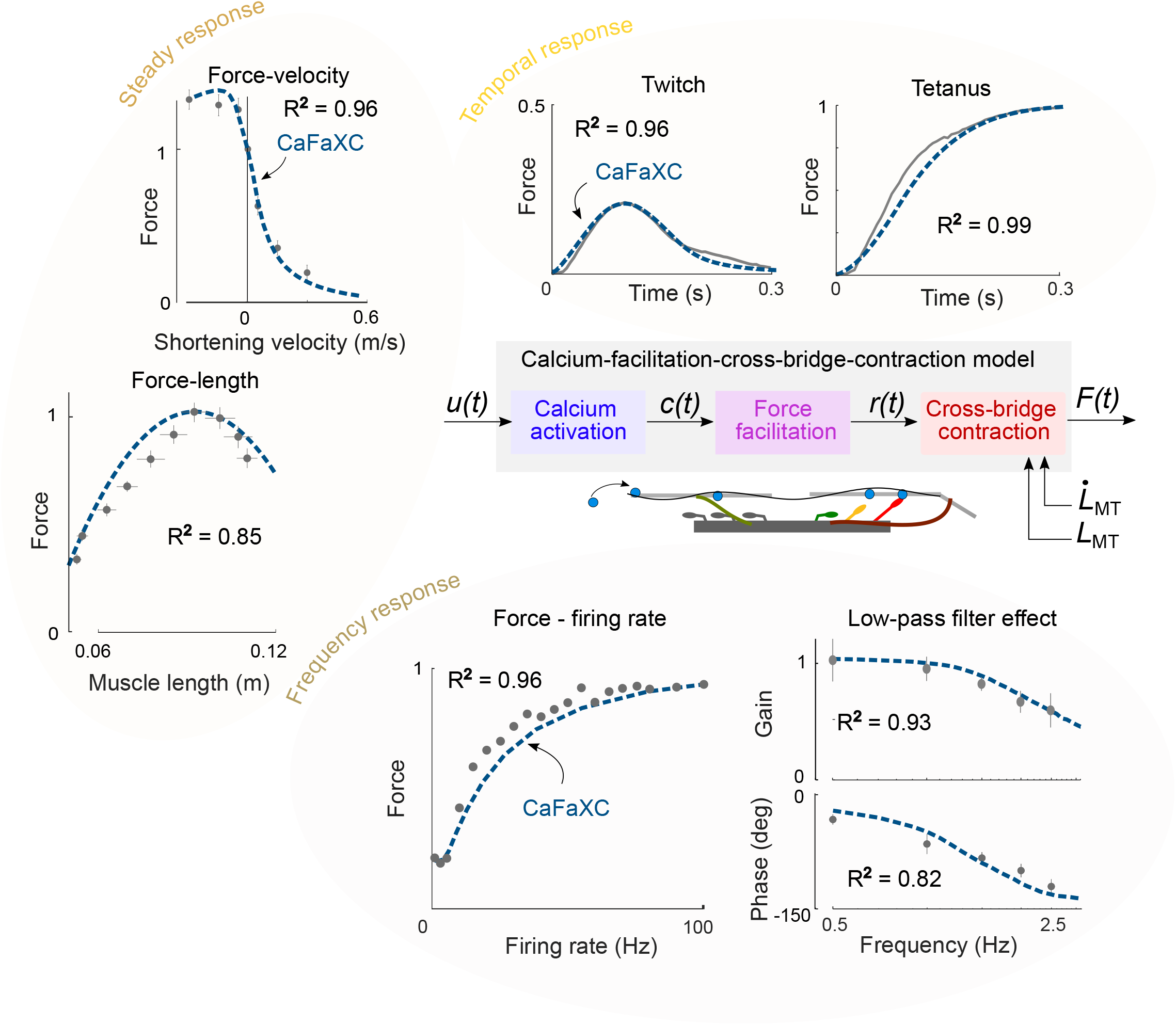
Proposed integrated muscle model and predictions for well-known responses of human quadriceps. (Center) Model integrates activation dynamics, facilitation dynamics and crossbridge dynamics with length feedback. Facilitation dynamics reflects processes intermediate to activation and crossbridge cycling. (Top left) Steady force-length and force-velocity responses of muscle contractile element, reproduced by proposed model (R^2^ = 0.85, and R^2^ = 0.96). (Top right) Temporal twitch and tetanus responses of muscle, reproduced well by proposed model (R^2^ = 0.96 and R^2^ = 0.99). (Bottom) Frequency responses of muscle, including force – firing rate and low-pass filter effect. Force-firing rate: force increases with stimulate rate, captured by proposed model (R^2^ = 0.96). Low-pass filter effect: Force gain, quantified as the amount of force per unit electromyography (EMG), decreases with cyclic contraction frequency, captured by proposed model (R^2^ = 0.93). The force phase, quantified as phase delay between EMG and force, increases in magnitude with cyclic contraction frequency, captured by proposed model (R^2^ = 0.82). All data are for human quadriceps *in vivo* (see Supplementary Materials and Methods) [force-length (Austin et al., 2010), force-velocity (Westing et al., 1990), twitch and tetanus (Verges et al., 2009), low-pass filter effect (van der Zee and Kuo, 2021), force-firing rate (Binder-Macleod and McDermond, 1992)]. Parameter values: τ _act_ = 2 ms, τ _deact_ = 50 ms, τ _fac_ = 60 ms, τ _def_ = 20 ms, *f*_1_ = 140 s^-1^, *g*_1_= 140 s^-1^, *g*_2_= 1388 s^-1^, *g*_3_= 78 s^-1^ (see Appendix for derivation and definitions). Lacking data from individual participants for some conditions, all reported R^2^ values are with respect to empirical average data.

## Discussion

This study aimed to quantitatively determine which dynamics underly muscle force development. Relatively slow dynamics are apparent from muscle’s temporal and frequency responses, but are not captured by conventional muscle models. We found that an additional process, termed force facilitation, is needed to reconcile slow force development with fast calcium activation. We next examine the relative timing contributions from our integrated model, consider mechanisms underlying the proposed force facilitation dynamics, and discuss the biological relevance and implications for movements.

These results resolve long-standing questions regarding activation dynamics. We found that any first-order definition of activation, whether coupled to Hill-type or to crossbridge mechanics, can only reproduce one of either tetanic activation or force development (Fig. 4), or one of tetanic and twitch forces (Figs. 4-5). With calcium imaging (Baylor and Hollingworth, 2003; Hollingworth et al., 1996; Zhao et al., 1996), activation can be defined specifically, for example as CaTrop here, or as free or total calcium concentration. But to agree with such data, a calcium-based definition requires fast activation dynamics. We found a time constant (2 ms) to be compatible with both human and mouse data, in agreement with fast calcium dynamics previously reported across a range of species and muscles (Baylor and Hollingworth, 2003; Rome et al., 1996). Such fast calcium dynamics therefore can account for only 3% to 5% of overall force development time. This contrasts with previous models that have qualitatively attributed activation dynamics to calcium processes (e.g., van Soest et al., 1993; Thelen, 2003; Mayfield et al., 2023). Both data and model now show that calcium dynamics, while critical to muscle function, are too fast to account for actual force development.

Additional dynamics are necessary to explain slow force development in muscles. In most biomechanics studies, muscle dynamics have comprised some subset of calcium-based activation, crossbridge mechanics, and Hill-type contraction against series elasticity and body. These dynamics can be resolved by applying empirical data and enforcing dynamical constraints. For example, Huxley’s (1957) crossbridge mechanics both explain and are constrained by the muscle force-velocity curve. This constraint required quite fast rate constants here, so that crossbridge dynamics accounted for only about 3% of force development time. Hill-type contraction dynamics also contribute to force development due to the force-velocity curve, as muscle fibers shorten against series elasticity (despite isometric conditions for the limb). Constrained by experimental data for human quadriceps, such contraction dynamics accounted for a substantial 27% to overall force development time. However, even the combination of calcium activation with conventional Hill-type and Huxley-type contraction dynamics can only explain about one-third of overall force development time.

We have therefore introduced a new latent state, provisionally termed force facilitation, to account for the missing force development time. Specifically, a first-order process following calcium dynamics is sufficient for compatibility with both calcium and force data. It presently lumps together more mechanistic processes (Fig. 6C) with a single time constant empirically determined to be quite slow, accounting for most (about 67% for mouse and 77% for human) of force development time. The need for such dynamics may not have been appreciated previously, because few models have addressed calcium activation and force development directly, and have thus not exposed the temporal gap between the two. The proposed force facilitation is necessary to account for that gap, and appears to be the time-dominating process in muscle’s ability to produce force, also known as latent activation.

The integrated model also provides insight regarding the frequency response behavior of whole muscle (Fig. 1 and Fig. 8). Muscle has long been observed to resemble a low-pass filter of excitation (Partridge, 1965; Stein et al., 1972; Milner-Brown et al., 1973; Bawa and Stein, 1976; Crosby, 1978; Winter, 2009). For example, force magnitude per unit excitation (gain) decreases with contraction frequency, similar to a critically damped second-order system with a relatively slow natural frequency (e.g., 2 - 3 Hz; Bawa and Stein, 1976; Milner-Brown et al., 1973). Muscle is thus far slower than explained by calcium activation (Fig. 7) or latent activation dynamics of typical Hill-type models (e.g., Zajac, 1989; Thelen, 2003; Millard et al., 2013), but the discrepancy was unclear because frequency response methods and electromyography-to-force transfer functions (e.g., Calvert and Chapman, 1977; Lloyd and Besier, 2003; Buchanan et al., 2004; Lee et al., 2011; Lee et al., 2013) usually provide little physical mechanism or location for the identified time constants. Our model with more mechanistic calcium and contraction dynamics suggests that force facilitation may be the dominant (slowest) dynamics underlying the low-pass filter effect (Figure 7), followed by contraction dynamics.

Force facilitation dynamics may be relevant to modeling of human movement. Lacking rate-limiting force development dynamics, Hill-type models with ad hoc first-order activation are often too fast. In musculoskeletal simulation (e.g., OpenSim: Seth et al., 2011), such models have been observed to underestimate the amplitude of muscle excitation transients for faster walking speeds (Luis et al., 2022), and EMG-to-force delays for walking (Luis et al., 2022) and hopping (Jessup et al., 2023). Such models also underestimate the energetic cost of faster movements, for example only predicting 7% and 40% of the cost increases for cyclic torque production (van der Zee and Kuo, 2021) and reaching movements (Wong et al., 2021). Other models have used slower activation time constants that agree better with (tetanic) force development data (Curtin et al., 1998; Lemaire et al., 2016). But such models may underestimate the amplitude of twitch forces (see Fig. 4-5), or fail to represent higher order dynamics evident in reaching movements (Murtola and Richards, 2023). In addition, attribution of timing to calcium rather than force facilitation dynamics could lead to incorrect estimates of calcium transport and its energetic cost (Szentesi et al., 2001; Barclay et al., 2007), potentially leading to underestimated energetics as stated above. The inclusion of force facilitation dynamics, whether in the present model or otherwise-conventional Hill-type models, may improve how well musculoskeletal simulations can perform when tested against independent experimental data. Such models can potentially improve upon predictions of muscle excitations, forces, and energetics during movement.

The present model also integrates together previously disparate dynamics. To date, Hill-type models have generally included dynamical interactions with body and series elasticity but not crossbridge dynamics, and crossbridge models the converse. Their combination has previously been found to be computationally impractical (van Soest et al., 2019), which we addressed by adopting the distribution moment approach (Zahalak, 1981), resulting in modestly greater complexity than Hill-type models (see Appendix). We found both types of dynamics to be relatively fast but nonetheless important for modeling movement. Crossbridges are the primary explanation for the force-velocity relation, which must arise from the same mechanics that govern force production. Such mechanics also predict transient muscle phenomena such as short-range-stiffness (Campbell and Lakie, 1998), observable in upright balance (De Groote et al., 2017) and implicated in proprioception (Simha and Ting, 2024). Crossbridge cycling also mechanistically predicts the energetics of contraction (Ma and Zahalak, 1991). This contrasts with Hill-type models that, due to phenomenological curve fits, enforce no mechanistic consistency (e.g. between force production and force-velocity) and have no mechanism for short-range stiffness. Such models are thus susceptible to overfitting (e.g., Figs. 4 and 5), making it difficult to make generalizing predictions. Here we guarded against overfitting by using twitch, tetanus, and/or calcium data to identify force facilitation, and then independently testing both the low-pass filter effect and the force-firing rate relationship (Fig. 8). This facilitates the combination of both crossbridge dynamics for mechanistic consistency, and a dynamical interface with the body derived from the Hill-type approach.

Although we advocate for force facilitation dynamics, its underlying mechanisms remain unclear. Potential processes include cooperative interaction between thin and thick filaments (Campbell, 2014; Walcott, 2014) and facilitative recruitment of myosin heads (Reconditi et al., 2011; McNamara et al., 2014; Linari et al., 2015; Irving, 2017; Nag and Trivedi, 2021; Ma et al., 2023), mediated by other protein interactions (Song et al., 2021; Squarci et al., 2023; Hessel et al., 2024). We provisionally lumped an overall effect into the force facilitation state *r*, identifying its time constant from force development data. It is thus presently the least mechanistic feature of our model, and is a latent state. This might seem like little improvement over Hill’s (1949) original active state, but force facilitation is more easily identifiable from experiment (e.g., Fig. 7), and more clearly localized to the processes between calcium activation and actual crossbridge binding. It suggests that Hill’s active state might be mechanistically interpretable as a measure of crossbridge binding, such as a fraction of attached crossbridges. Further experimentation and modeling is needed resolve how muscle force is actually facilitated (e.g., Longyear et al., 2017; Campbell et al., 2018;Liu et al., 2024; Squarci and Campbell, 2024). There is a need to assess and compare force facilitation across slow to fast muscle fiber types, between species, and within a species but between muscles. But in contrast to the Hill-type models that also reproduce forces matching data (Curtin et al., 1998), force facilitation here highlights the timing gap between calcium activation and force development.

Additional limitations of the CaFaXC model include the need for two additional model parameters compared to Hill-type models. The model’s contraction dynamics parameters may be mapped directly from Hill-type models (see Appendix), but the separate calcium and force facilitation dynamics requires two additional parameters, for forward and backward calcium-based activation rates, distinct from current Hill-type models. These parameters are identifiable from in vitro CaTrop estimates (Fig. 7A) where available. But for human, we used (*in vivo* electrically-stimulated) force responses (Fig. 7B) to provisionally identify both activation and force facilitation parameters simultaneously. This experimental procedure is in principle applicable to a variety of human muscles, facilitating quantitative determination of force development that contrasts with the ad hoc parameters used in many Hill-type models. However, a disadvantage is that the model’s calcium activation parameters have yet to be tested against independent measures of CaTrop for human, presently unavailable. Such data would also be helpful for testing more complex models of calcium transport and binding (e.g., Baylor and Hollingworth, 2003; Kim et al., 2015). We consider the CaFaXC model to represent the minimum necessary to reproduce both calcium activation and force development data while having identifiable parameters.

We have thus far used relatively simple crossbridge dynamics, as in Huxley’s classic model (Huxley, 1957). While more complex crossbridge models have become available (Campbell and Lakie, 1998; Campbell and Moss, 2002; Campbell, 2014; Walcott, 2014; Campbell et al., 2018; Kosta et al., 2022; Liu et al., 2024), the classic model has remained valuable (Lemaire et al., 2016; Newhard et al., 2019; van Soest et al., 2019), in part because it mechanistically explains force-velocity and energy-velocity relationships (Huxley, 1957). These relationships serve as constraints on the model, which must also explain isometric force and force development transients and temporal and frequency response dynamics (Fig. 8). This requires only a small number of parameters and helps guard against model over-fitting. While other, more complex, crossbridge models are valuable for detailing features such as weakly-bound crossbridge states (Huxley and Simmons, 1971; Jarvis et al., 2021), and force transients during lengthening (Lombardi and Piazzesi, 1990), we found the classic crossbridge model sufficient to test force development dynamics, and consider it adaptable to more complex models.

The present model is amenable to additional features such as force transmission and energetics. Musculoskeletal models often include muscle pennation as part of force transmission to the skeleton (Thelen, 2003; Millard et al., 2013; D’Hondt et al., 2024), and some also estimate metabolic energy expenditure (e.g., Umberger et al., 2003), albeit phenomenologically. The present model’s contractile apparatus has the same external interface as Hill-type models, and is thus readily adaptable to pennation. Owing to the DM approximation, CaFaXC’s modest computational cost (about twice that of the Hill-type model, see Appendix) facilitates implementation in movement simulation. The model is also adaptable to energetics, because it includes crossbridge binding and calcium transport, which are mechanistically implicated in energetics (Szentesi et al., 2001; Barclay et al., 2007). It remains to be tested whether the CaFaXC model predicts the energetics better than Hill-type models, but we consider the mechanistic approach to have greater potential, particularly because most muscle experiments are aimed at mechanistic understanding rather than phenomenological curve fitting.

## Conclusions

The rate-limiting dynamics of force development are due to neither calcium activation, nor crossbridge cycling, nor shortening against elastic tissues, which are all too fast to explain data. Rather, an intermediate set of “force facilitation” dynamics appear necessary to explain time- and frequency-dependent aspects of muscle force development in a self-consistent manner. A model of force facilitation introduces a relatively slow, dynamic relation between muscle excitation and force, with potential applications to predicting the mechanics and energetics of movements.

## List of symbols and abbreviations

*L*_*M*_: Muscle fiber length
*LT*: Tendon length
*L*_*MT*_: Muscle-tendon complex length (i.e. *L*_*M* +_ *L*_*T*_)
*F*_*CE*_: Contractile element force
*F*_*PE*_: Parallel element force
*F*: Muscle force (i.e., *F*_*CE*_+ *F*_*PE*_)
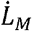: Muscle fiber velocity
*a*: Muscle activation
*c*: Calcium activation
*r*: Force facilitation
*u*: Muscle excitation
*k*_*cE*_: Contractile element stiffness
*n*(*x*): Distribution of attached crossbridges
*Q*_i_(*x*): i-th order moment of the distribution of attached crossbridges
*x*: Crossbridge strain
τ_a_: Activation dynamics time constant
τ_act_: Activation dynamics forward time constant
τ_deact_: Activation dynamics backward time constant
τ_r_: Facilitation dynamics time constant
τ _fac_: Facilitation dynamics forward time constant
τ_def_: Facilitation dynamics backward time constant
*f*: Crossbridge cycling attachment function
*f*_1_: Crossbridge cycling detachment function
*g*_1_: Crossbridge attachment rate constant (within attachment zone) *“*_1_ Crossbridge detachment rate constant within attachment zone
*g*_2_: Crossbridge detachment rate constant outside attachment zone and negative strain
*g*_3_: Crossbridge detachment rate constant outside attachment zone and positive strain

## Appendix

### Steady model properties for human quadriceps muscle *in vivo*

We modelled the human quadriceps as a single muscle-tendon complex, lumping properties of the four individual heads together. Parameter values were either obtained from previous models, or derived from previously reported data (see Table S1).

#### Contractile element

The contractile element force-length relation was assumed to be a quadratic function of muscle fiber length *1*_L_ *L*_M_ (van Soest and Bobbert, 1993). Height, width and horizontal location of this force-length relation are specified by three parameters (i.e., _max_, *‘“*, and *L*_M,opt_ respectively, see Table S1). For the Hill-type model, we employed a standard force-velocity relation (Umberger et al., 2003) specified by maximal shortening velocity, curvature and eccentric force asymptote (*v*_max_, *A*,_rel_ and *F*_asym_ respectively, see Table S1). Parameter values were chosen to approximately agree with torque-angular velocity data for human knee extension (Westing et al., 1990). For the crossbridge model and CaFaXC model, we fitted attachment and detachment rate constants to yield the Hill-type force-velocity relation in steady state conditions (see “Parameter estimation - Crossbridge rates”).

#### Series elastic element

The series elastic element force was assumed to increase linearly with strain for large forces, with exponential toe-region at small forces as described previously (Winters, 1995). Given the estimated tendon slack length *L*_*T*0_ (van Soest et al., 1993), we chose the remaining parameters (i.e.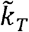, *ρ*_*toe*_, *ϵ*_*toe*_, *σ*_*toe*_) to agree with previously reported ultrasonography-based series elastic length changes in human quadriceps (van der Zee and Kuo, 2021).

#### Parallel elastic element

The parallel elastic element force was assumed to increase quadratically with strain above passive element slack length *L*_M,0_, and to reach maximal isometric force *F*_max_ at a specified strain ϵ_*M*_ as described previously (van Soest et al., 1993), with parameter values from literature (Thelen, 2003).

### Dynamic model properties for human quadriceps muscle *in vivo*

#### Activation dynamics

We used a common first-order formulation for all models:

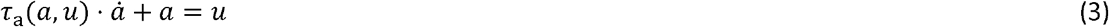

Activation time constant τ_a_ equals. τ_act_ when excitation exceeds activation and equals. τ_deact_ otherwise. In the proposed CaFaXC model, activation is explicitly defined as CaTrop. The same formulation as in Eqn 3 is applied, but activation *a* is replaced with calcium activation *c*. Time constants were fit to agree with either calcium activation or force development (see “Parameter estimation – Time constants”).

#### Force facilitation dynamics

We used a first-order formulation of dynamics between calcium activation *c* and force facilitation *r*:

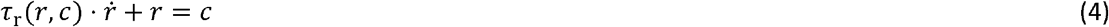

As multiple mechanisms may underly force facilitation dynamics, the forward and backward dynamics may be different. Like activation dynamics, facilitation time constant τ_r_ therefore equals. τ_fac_ when activation exceeds facilitation and equals. τ_def_ otherwise. Like activation dynamics, time constants were fit on twitch and tetanus force development (see “Parameter estimation – Time constants”).

#### Crossbridge model rate functions

The crossbridge model behavior depends on the attachment and detachment rate functions *f*(*x*) and *g*(*x*). In line with a previous model formulation (Zahalak, 1981):

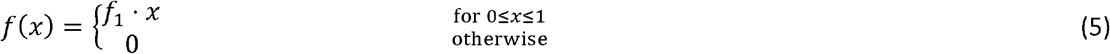

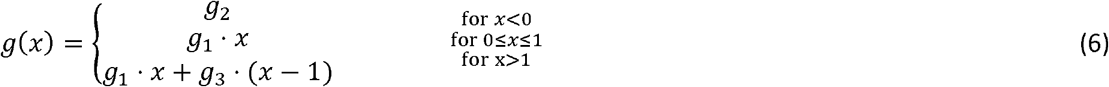

Here, crossbridge strain *x* is normalized for the reach of the myosin molecule *h*, which we assume to be 12 10^−9^ m in line with previous models (Newhard et al., 2019) and experimental data (Finer et al., 1994). To reflect activation-dependent changes in *v*_max_ (Chow and Darling, 1999), all rate constants were linearly scaled with activation *r* (crossbridge model) or force facilitation *r* (CaFaXC). Specifically, rates ranged between their maximum value (i.e., *f*_1_, *g*_0_,*g*_1_,*g*_2_) in maximally active muscle (e.g.r =1)and a fraction *μ* of that maximal value in inactive muscle (e.g., *r= 0*. We chose 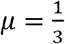 for human quadriceps, similar to the *v*_max_ - scaling employed in the Hill-type model force-velocity relation (Umberger et al.,2003).

### Geometrical scaling of crossbridge dynamics

When incorporating crossbridge dynamics into standard muscle modelling geometry (Fig. 2A), one needs to assure that the forces in the three elements (i.e., CE, PE, SE) are compatible. This is trivial in Hill-type models, because CE force *F*_*CE*_ is determined from forces in the elastic elements (i.e., *F*_*PE*_ and *F*_*SE*_). This is not trivial in crossbridge models, because *F*_*CE*_ must simultaneously be compatible with forces in elastic elements and with contractile element crossbridge dynamics. As conceived previously (Stoecker et al., 2009; Lemaire et al., 2016), this was accomplished through applying a constraint on the time-derivatives of the forces (i.e.,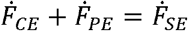) at each integration step.

The first-order moment of the crossbridge distribution *n* equals the crossbridge force 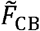, expressed relative to the force when all crossbridges are attached at uniform strain within the attachment range 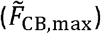. The resultant force of the crossbridge model contractile element *F*_*CE*_ depends on the total number of crossbridges and the crossbridge stiffness, which are both difficult to estimate *in vivo*. We therefore employed a force scaling parameter *δ*_*F*_ that assures that the crossbridge model contractile element force equals that of the Hill-type model in isometric conditions, similar to a previous model (Lemaire et al., 2016):

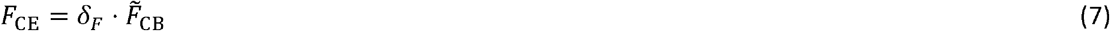

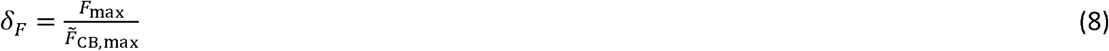

Here, *F*_max_ is the maximal isometric force of the contractile element, as in the Hill-type model. For non-isometric contraction, muscle fiber contraction velocity 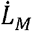 relates to crossbridge strain rate 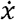 :

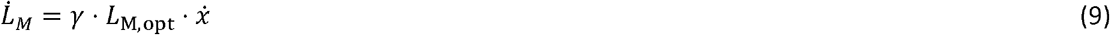

The scaling parameter *γ* is the gear ratio between crossbridge strain *x* and half-sarcomere displacement:

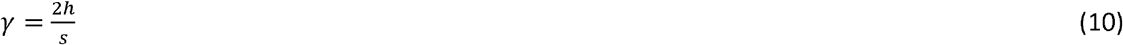

Here *s* is the sarcomere length at maximal filament overlap, which took to be 2.64 10^−6^ m for human muscle (Walker and Schrodt, 1974), resulting in a *γ* of 0.009.

### Cross-bridge dynamics and distribution-moment approximation

Conventional crossbridge model dynamics are described by the following partial differential equation:

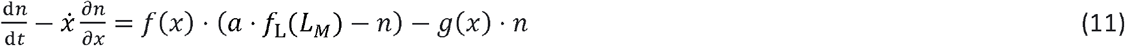

where *n* is the crossbridge distribution, *f*(*x*) and *g*(*x*) are the rate functions (Huxley, 1957), and *“* is the muscle activation (replaced by *r* in CaFaXC) and *f*_L_ (*L*_M_) is the force-length overlap (e.g., Lemaire et al.,2016). This partial differential equation can be transformed into a set of ordinary differential equations by method of characteristics (Zahalak, 1981), which can be time-integrated numerically (Lemaire et al., 2016). This incurs a high computational cost because (1) the discontinuous nature of the rate functions (Eqns 5-6) results in a stiff equation, (2) the strain vector *x* needs to be spaced densely and have a broad range. The high stiffness of the equation means that the integration step size must be small, while the strain vector constraints require a large number of states. In practice, about 18000 states are needed (van Soest et al., 2019). This is because keeping isometric force errors below 0.5% requires 200 bins between *x* = 0 and *x=*1, and whole-muscle length changes during movement can be similar to *L*_M,opt_, equivalent to strain changes *Δx* of 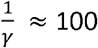. The overall number of strain bins equals the product of bin density (200) and strain range (100), for a total of about 20000.

As conceived by Zahalak (1981), *n*(*x*) may be approximated by a Gaussian and the relevant mechanics expressed in terms of its first three moments *Q*_0 ™_*Q*_2_. For maximal activation, Eqn 11 can be rewritten as a function of the λ-th order moments of the distribution *Q*_*λ*_:

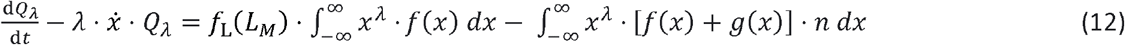

Zahalak and colleagues later adapted Eqn 12 for submaximal activation (Zahalak and Ma, 1990; Zahalak and Motabarzadeh, 1997), by scaling the attachment rate function *f* (*x*) with activation *a* However, when simulating whole muscle as opposed to single fibers, Lemaire et al. (2016) noted that it may be more appropriate for activation to set the total available crossbridges. We therefore applied the form of Lemaire et al. (2016) to adapt Eqn 12, but using facilitation *r* instead of activation *a*:

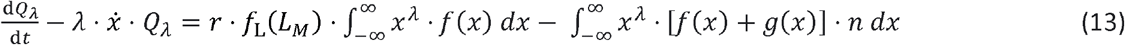

The DM model thereby only requires three states to describe the crossbridge distribution during contractions, much less than the 18000 states of the original model. For certain rate functions *f*(*x*) and *g*(*x*) including the piece-wise linear functions employed here (see Eqns 5-6), the associated spatial integrals can be computed analytically (Zahalak, 1981). Furthermore, the differential equation is much less stiff, because the discontinuous rate functions are smoothened through spatial integration. The combination of larger integration steps and fewer states result in a 700 times smaller computational cost of the CaFaXC model compared to the conventional crossbridge model for the tetanus contractions of human quadriceps simulated here (0.08 s vs. 56 s on a modern laptop, Intel Core i7-10850H @ 2.70GHz, using MATLAB’s ode113 with a variable integration step size smaller than 0.001 s), while yielding similar muscle forces (see Figs. S1-S2). The CaFaXC model is about 2 times slower than the Hill-type model with force-based activation dynamics.

### Parameter estimation

#### Crossbridge cycling rates

The crossbridge cycling rates uniquely determine the crossbridge model’s steady-state force-velocity relation. As conceived previously (Huxley, 1957; Newhard et al., 2019), the shortening aspect of the crossbridge model force-velocity relation is described by:

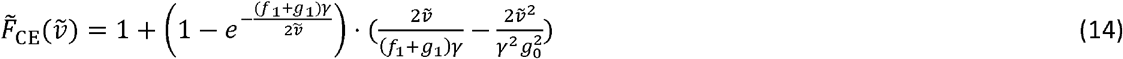

The lengthening aspect of the relation is described by:

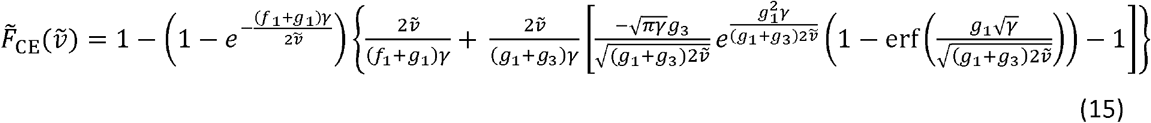

Here, velocity 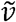 is the relative sliding velocity, as expressed in optimum lengths (e.g. 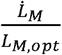). Given the geometric scaling parameter *γ* and constraining the detachment rate within the attachment zone to equal the attachment rate (i.e. *g*_1_=*f*_1_), we chose the remaining rate constants such to minimize the squared difference between the crossbridge model and the Hill-type model force-velocity relations (MATLAB’s fminsearch). Crossbridge cycling rate constants are reported in main text (Figure legends).

#### Time constants

As described in main text, the activation dynamics time constants of the conventional muscle models were fit to approximately agree with either empirical calcium activation or force development. The proposed CaFaXC model has one additional set of time constants, for a total of four time constants. These time constants were chosen to best match four features of force development: (1) tetanus rise time (τ_act_,. τ_fac_), (2) twitch (or tetanus) decay time (τ_deact_,. τ_defac_), (3) twitch time-to-peak (.τ_act_,. τ_deact_,. τ_fac_) and (4) twitch amplitude (.τ_act_,. τ_deact_,. τ_fac_). The additional experimental requirement for parameter estimation of the proposed model compared to conventional models is therefore assessment of twitch forces. Fortunately, twitch and tetanus require very similar experimental methods, and typically only differ in the duration of stimulation (brief or long). Here, we chose these time constants to approximately agree with twitch and tetanic force development for human quadriceps, while also evaluating force development in other conditions (e.g., force – firing rate, force – contraction frequency). The values for the activation dynamics time constants are in line with previous estimates of active state rise and decay in human muscle (Hof and Van den Berg, 1981b). Time constants are reported in main text (Figure legends).

### Parameter values for mouse extensor digitorum longus muscle *in vitro*

Parameter values for mouse extensor digitorum longus (EDL) fast-twitch fibers were estimated for the *in vitro* conditions considered here (Hollingworth et al., 1996). Hollingworth and colleagues estimated that 1-10 fibers were producing force, and that the average fiber diameter was 43 *μ*m. Assuming an average of 6 isolated fibers, and a maximal tetanic tension of 390 kN/m^2^ (Lännergren and Westerblad, 1987), maximal force of this fiber bundle was estimated at 0.0017 N. Muscle optimum length *L*_M,opt_ was estimated at 0.014 m (Hakim et al., 2013) and all other length parameters were assumed to scale linearly with the human quadriceps values. Maximal shortening velocity *ν*_max_ was estimated at 10 *L*_M,opt_ *s*^-1^, considering that: (1) mouse EDL is believed to be uniformly fast-twitch (Hollingworth et al., 1996), (2) smaller mammals typically have faster maximal shortening velocities than larger mammals (Wakeling et al., 2012). Because activation-dependent *v*_max_-scaling is thought to arise from recruitment of faster motor units (Petrofsky and Phillips, 1981), we did not apply this to the fast-twitch mouse muscle fibers (i.e. *μ=1*). Force-velocity curvature *A*_*rel*_ was estimated at 0.3, deemed appropriate for fast-twitch fibers (Wakeling et al., 2012). As with the human quadriceps, mouse EDL crossbridge cycling rates were chosen as to fit the force-velocity relation (MATLAB’s fminsearch). The same value of *h* was used for crossbridge scaling, but a smaller sarcomere length of 2.4 10^−6^ m (Goulding et al., 1997) yielded a slightly larger geometric scaling parameter *γ* (about 0.01).

## Competing interest

The authors declare no competing or financial interests.

## Author contributions

Conceptualization: T.J.v.d.Z., A.D.K.; Methodology: T.J.v.d.Z., A.D.K., J.D.W.; Software: T.J.v.d.Z., J.D.W.; Validation: T.J.v.d.Z., J.D.W.; Formal analysis: T.J.v.d.Z.; Investigation: T.J.v.d.Z.; Resources: A.D.K.; Data curation: T.J.v.d.Z.; Writing - original draft: T.J.v.d.Z.; Writing - review & editing: T.J.v.d.Z., A.D.K., J.D.W.; Visualization: T.J.v.d.Z., A.D.K.; Supervision: A.D.K.; Project administration: A.D.K.; Funding acquisition: A.D.K.

## Funding

This work was supported by the Natural Sciences and Engineering Research Council of Canada (NSERC Discovery and Canada Research Chair, Tier 1).

## Data availability

All data and code are available at github.com/timvanderzee/CaFaXC.

## Notes

### Competing Interest Statement

The authors have declared no competing interest.

